# Gametic Selection and Mating Systems show Mutually Dependent Evolution

**DOI:** 10.1101/2023.02.08.527794

**Authors:** Michael F Scott, Simone Immler

## Abstract

Competition among pollen or sperm (gametic selection) can cause evolution. Mating systems shape the intensity of gametic selection by determining the competitors involved, which can in turn cause mating system evolution. We model the bidirectional relationship between gametic selection and mating systems, focussing on variation in female mating frequency (monandry-polyandry) and self-fertilisation (selfing-outcrossing). First, we find that mating systems affect evolutionary responses to gametic selection, with more effective gametic selection when fertilisation success depends on haploid genotypes, rather than the diploid genotype of the father. Monandry and selfing both reduce the efficacy of gametic selection despite creating intense selection among pollen/sperm from heterozygous males with haploid expression. This effect means selfing can increase mutation load, in contrast to classic predictions where selfing purges deleterious mutations. Second, we show that mating systems can evolve via their effect on gametic selection, with polyandry evolving because it removes deleterious alleles more efficiently and increases offspring fitness. Our population genetic models reveal that this ‘good sperm’ effect could plausibly give a selective advantage for polyandry over monandry of only around 1%. Selection for polyandry is lessened further if some loci experience balancing selection and is likely to be overwhelmed by any direct fitness effects of mating systems. Similarly, the indirect benefits from manipulating gametic selection have a weak influence on the evolution of selfing, which is dominated by ‘automatic selection’ and inbreeding depression in our model. Nevertheless, gametic selection can be potentially decisive for selfing evolution because it significantly reduces inbreeding depression, which favours selfing. One test of the predicted interactions between gametic selection and mating system evolution would be to compare evolutionary rates in genes with different expression patterns across different mating systems.

## Introduction

Males typically produce a large number of gametes or gametophytes (hereafter male gametes) that then compete to fertilise a small number of eggs or ovules (Trivers 1972) generating considerable selective pressure (hereafter gametic selection). The pool of male gametes that compete against one another depends on ‘who mates with whom’, which is called the mating system. Two important aspects of mating system variation are the number of males that females mate with (Kokko et al. 2014) and the rate of self fertilisation (Barrett 2014). That is, mating systems vary from monandry (females mate with one male) to polyandry (females mate with several males) and from selfing (where male and female gametes are derived from one individual) to outcrossing (where male and female gametes are derived from different individuals). Both axes of mating system variation affect the genetic composition of male gamete pools and with that gametic selection. We model interactions between mating systems and gametic selection from two angles, one to study evolutionary responses to gametic selection under different mating systems and one to study the evolution of mating systems with gametic selection.

Evolutionary responses to gametic selection depend on the way genetic material is expressed, with significant variation across genes and taxa. Fertilisation success may depend on a gamete’s haploid genotype or the diploid genotype of the male that produced them. In plants, fertilisation success is typically thought to depend on the gametophyte’s own haploid genotype (Mulcahy et al. 1996, Tonnabel et al. 2021) because pollen tubes are multicellular and express 60-70% of all genes (Borg et al. 2009, Qin et al. 2009). Indeed, haploid expression and pollen competition has been shown to cause non-random inheritance of genotypes from a single male (Stehlik and Barrett 2006, Leppälä et al. 2013, Williams and Mazer 2016, Swanson et al. 2016, Corbett-Detig et al. 2019) and pollen-expressed genes show stronger signatures of selection than random genes (Arunkumar et al. 2013). In animals, on the other hand, success during sperm competition is usually assumed to depend on the father’s diploid genotype (Parker 1970). This assumption is based on the cytoplasmic bridges that link developing spermatids, allowing transcript sharing and effectively diploid expression at most genes (Joseph and Kirkpatrick 2004). Nevertheless, recent results suggest that haploid expression and selection in animal sperm has been underestimated (reviewed in Immler and Otto 2018, Immler 2019, Kekäläinen 2022). For example, sperm selection assays within single ejaculates of the zebrafish *Danio rerio* have been shown to cause allelic biases (Alavioon et al. 2017). Single cell expression data from primate testes has revealed extensive expression at late stages of spermatogenesis, with these genes experiencing accelerated evolutionary rates (Murat et al. 2022). Single cell expression is biased towards a haploid allele at 31-52% of spermatid-expressed genes in a range of mammals (Bhutani et al. 2021), approximately 20% of all genes. These results suggest that expression of different genes in animal sperm can vary continuously from haploid to diploid depending on degree of allelic bias (Navarro-Costa et al. 2020). Our models allow a range of allelic bias scenarios to allow the effect of gametic selection to be compared across genes and taxa.

It is not straightforward to predict how mating systems and gametic selection interact to produce evolutionary responses. First, single males produce gamete genotypes in equal proportions, which maximises the response to selection at heterozygous loci with haploid expression. Therefore, it is plausible that monandrous mating can increase gametic selection with haploid expression despite eliminating gametic selection with diploid expression. Second, haploid expression and selfing both affect the efficiency of purifying selection and the associated mutation load. In diploid heterozygotes, a homologous gene copy can (partially) mask a deleterious allele’s effect, preventing them from being efficiently removed by selection (Crow and Kimura 1965, Kondrashov and Crow 1991). Haploid expression means alleles are exposed to selection and selfing reduces masking by increasing homozygosity so both can reduce mutation load (Crow and Kimura 1965, Charlesworth et al. 1990, Charlesworth 2006). However, this effect is reversed when some alleles are favoured during gametic selection but reduce the fitness of diploid adults (Walsh and Charlesworth 1992, Immler et al. 2012, Otto et al. 2015). Under such a scenario, adult fitness would be optimised with less gametic selection.

The evolutionary responses to the interaction between mating systems and gametic selection is therefore rather complex and models are a useful way to examine these interactions. A previous model assuming haploid expression compared the expected mutation load between selfers and outcrossers (Charlesworth et al. 1990), and another theoretical study assessed the evolutionary outcome across selfing rates where gametes and adults have opposing selection pressures (Peters and Weis 2018). Most models of sperm competition assumed diploid control over sperm competition success, which means that gametic selection only occurs under polyandry (reviewed in Pitnick and Hosken 2010, Sutter and Immler 2020). Nevertheless, two theoretical studies have examined genes with haploid expression in sperm and no adult effect, finding that haploid expression can allow evolution under monandry (Ezawa and Innan 2013) and that evolutionary rates increase with haploid expression and the harmonic mean of the number of mates per female (Dapper and Wade 2016). The most relevant study to consider the evolution of mating systems via their effect on gametic selection examines the ‘good sperm’ hypothesis using a quantitative genetics framework (Yasui 1997), finding that polyandry can evolve if mutations reduce viability and viability is positively correlated with sperm competitiveness. In our population genetic approach, the correlation between viability and gametic selection arises from the allele frequency dynamics at each locus. One use of our model is to approximate the plausible selective advantage of polyandry via the ‘good sperm’ hypothesis in terms of standard population genetic parameters. Another advantage of our treatment is that we consider a range of mating systems and expression patterns in the same framework to offer comparative predictions for empirical testing.

We aim to model the two-way interaction between gametic selection and mating systems. We first consider the influence of mating systems (monandry/ polyandry and selfing/outcrossing) and gametic expression patterns (ranging from haploid to diploid) on the evolution of alleles that affect fertilisation success. We then allow the mating systems to evolve to examine how their effect on gametic selection influences mating system evolution. Despite having similar qualitative effects on the strength of gametic selection, we find that monandry and selfing are expected to follow distinct evolutionary trajectories.

## Model

We investigate both evolution under different mating systems and the evolution of mating systems themselves. We consider two mating system scenarios: (a) mating systems fall on a spectrum from polyandrous to monandrous and (b) mating systems can vary from selfing to outcrossing. We are specifically interested in the effects of competition among sperm or pollen.

Here, we outline the important features of our model. We provide a detailed model description in Appendix I and a supplementary data file that can be used to replicate the results. For simplicity, microgametophytes (pollen) and megagametophytes (inside the ovule) are called male and female gametes, respectively, and selection during competition among pollen or sperm is called gametic selection.

Primarily, we use a two-locus model with a fitness locus (**A**) that experiences selection directly and a modifier locus (**M**) that determines the mating system. The results from the two-locus model can then be extrapolated to multiple loci, see Appendix II. The alleles (*A* and *a*) at the fitness locus can have different effects on the fitness of adults of both sexes where fitness coefficients can differ between male and female adults and of male gametes; all the fitness terms are given in Table 1. In hermaphrodites, alleles can differentially affect male and female fecundity (Schärer et al. 2015) and therefore we also allow for separate male and female fitness effects in this scenario. We assume that gametic selection occurs among male gametes and that all female gametes are fertilized.

**Table 1:**
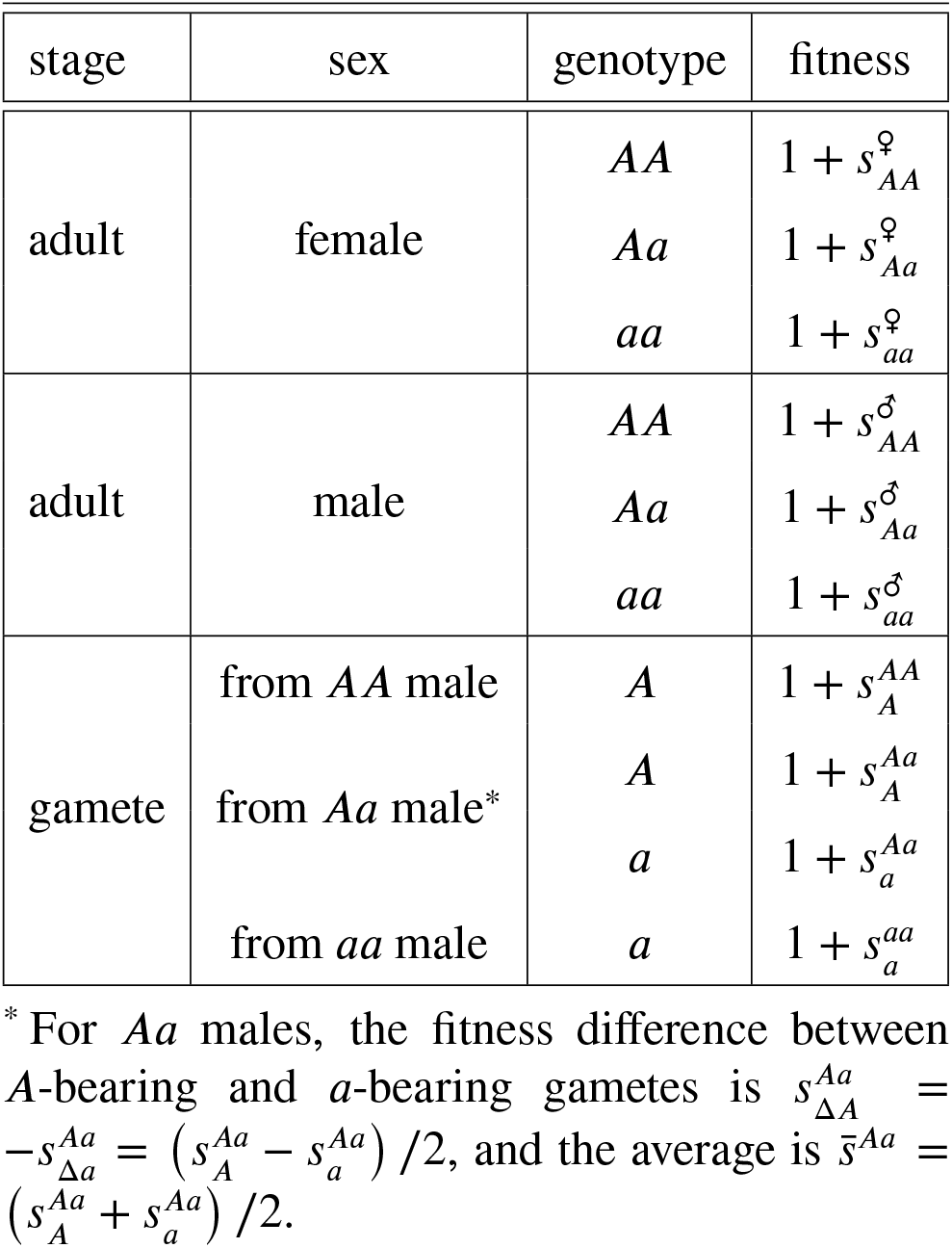
Fitness parameters for different life cycle stages.

Male gametic fitness depends on the expression of genetic material. Male gametes may express their own haploid genotype or they can express the diploid genotype of their father. For example, transcripts can be shared across cytoplasmic bridges between developing spermatids such that fully developed sperm express a mixture of the father’s alleles. We therefore allow male gametic expression to be haploid-like or diploid-like, or some combination. Because we allow male gametic expression to vary continuously from haploid-like to diploid-like, we use a continuous version of allelic dominance, as illustrated in Figure 1a.

**Figure 1:**
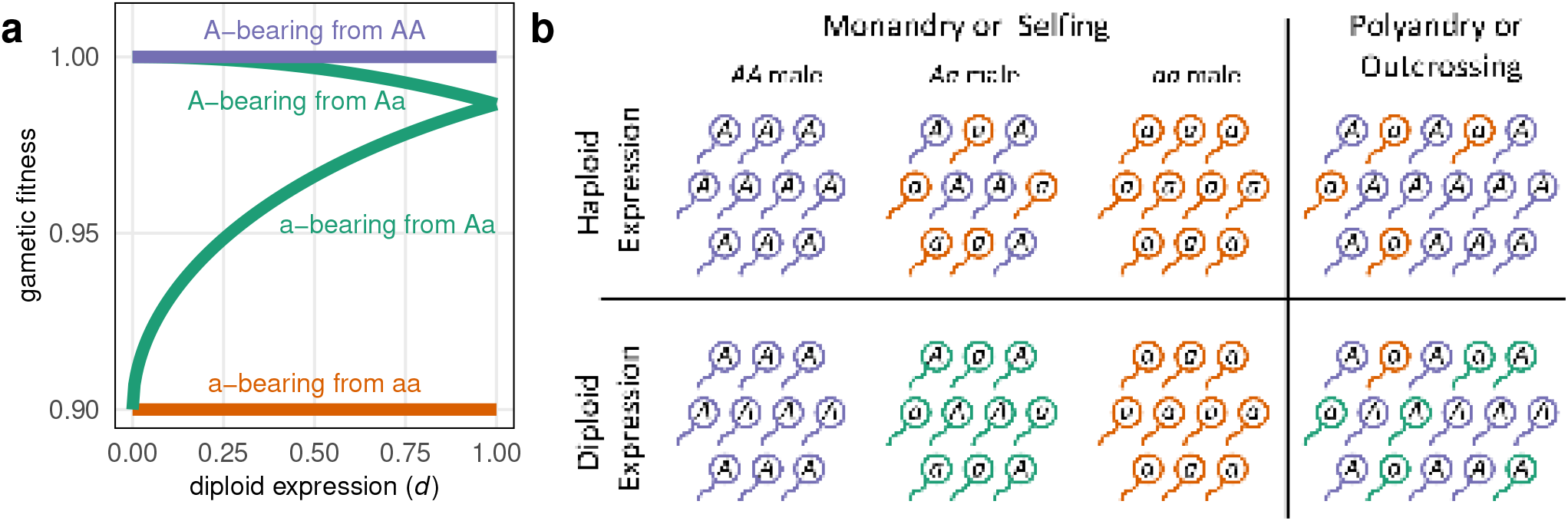
Illustration of competition among male gametes. (a) Gametic expression can vary continuously from fully haploid to fully diploid, with intermediate expression between. Here, the *a* allele is deleterious (*t* = 0.1) and partially recessive (*H* = 2). (b) Male gametes compete in local pools according to the mating system (colours represent gametic fitness). When expression is fully haploid, heterozygous males create highly competitive gamete pools where allele frequencies are equal, but there is no fitness variation with diploid expression. Monandry involves outcrossing between male and female individuals whereas selfing involves the male and female gametes from the same individual. The frequency of the different male gamete pools under monandry/selfing and the allele frequency in the polyandry/outcrossing gamete pool depends on the population genotype frequency.

The way male gametes compete among each other is determined by the mating system (Figure 1b). Mating systems can be a mixture of (i) polyandry and monandry or (ii) outcrossing and selfing. The mating system is controlled by the **M** locus genotype of the mother, which is initially assumed to be fixed for allele *M*. With a mixture of polyandry and monandry, a fraction (Π) of female gametes will be mated polyandrously (with many males competing for fertilisation) and the remaining fraction (1 –Π) is mated monandrously (with a single male). With a mixture of outcrossing and selfing, a fraction (Ω) of female gametes are outcrossed and the remaining fraction (1 – Ω) are selfed. The mating system is controlled by the **M** locus genotype of the female, which is initially assumed to be fixed for allele *M*. To examine mating system evolution, we introduce a new allele, *m*, that causes females to change their mating system allocation (to ⊓*_Mm_* or Ω*_Mm_* for *Mm* females and to ⊓*_mm_* or Ω*_mm_* for *mm* females).

Using male gametes for selfing may reduce the number of male gametes that are available for outcrossing; this is called ‘pollen discounting’ and has an important role in the evolution of selfing (Nagylaki 1976, Holsinger et al. 1984). Although we follow convention by using the term ‘pollen discounting’, our models are also applicable to hermaphroditic animals (e.g., Leonard 2018, Cutter 2019). In our model, pollen discounting is determined by *c*. When *c* = 0, selfing does not reduce the number of male gametes available for outcrossing. When *c* =1, increased selfing results in a proportional decrease in the number of male gametes that are available for outcrossing.

Monandry and selfing both create a similar selective arena for haploid selection (equation A2), in which gametes from only one male compete for fertilisation (Figure 1b). Polyandry and outcrossing create selective arenas where all male gametes in the population compete in a common pool (equation A3). However, under monogamy, mating occurs between different individuals (equation A4) whereas, under selfing, male gametes will fuse with female gametes produced by the same individual (equation A5). Thus, selfing and monandry have similar direct effects on the intensity of haploid selection but selfing will also increase homozygosity.

## Results

We assess evolution under different mating systems by looking at allele frequency trajectories at the fitness locus **A**. Initially, we assume all individuals have the same mating system by assuming they all carry the same modifier allele, *M*. We then allow the mating system to evolve by introducing a new modifier allele, *m*, that changes the mating system. To calculate these evolutionary trajectories, we assume that selection is weak (of order *ϵ* where *ϵ* ≪ 1). We further assume that the number of new mutations per locus per generation is very small (*μ* of order *ϵ*^3^). We extrapolate our results across multiple loci to estimate genome-wide mutation load and inbreeding depression. To do this, we assume that fitness effects across loci are uniform, multiplicative, and non-epistatic and that loci are loosely linked such that their frequencies can be considered independently (see Appendix II).

### Invasion and Fixation

We first derive deterministic invasion conditions at a single selected locus (**A**). If an allele’s frequency increases when it is rare, then it is able to “invade” and spread. If the allele frequency continuous to increase when it is at high frequency, then it is expected to reach “fixation”. As there are two alleles, fixation of one allele implies that the other allele does not invade. When both alleles can invade when they are rare, an intermediate equilibrium frequency is reached where both alleles are maintained by balancing selection.

Invasion by allele *n* (either allele *A* or allele *a, n* ∈ {*A, a*}), is determined by *I_n_*. When *I_n_* is positive (*I_n_* > 0), selection favours the spread of a rare *n* allele (e.g., a new mutant). We express *I_n_* in terms of the selection that occurs in females 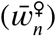 and selection in males, which is further divided into diploid male selection 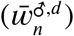 and male gametic selection 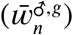 to give

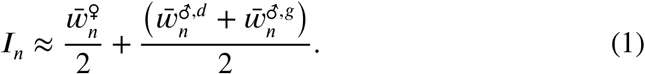

For different mating systems (polyandry/monandry or outcrossing/selfing) we give these selection terms in table 2. In short, *I_A_* gives the selective advantage of a rare *A* allele among predominantly *a* alleles (vice versa for *I_a_*).

**Table 2:**
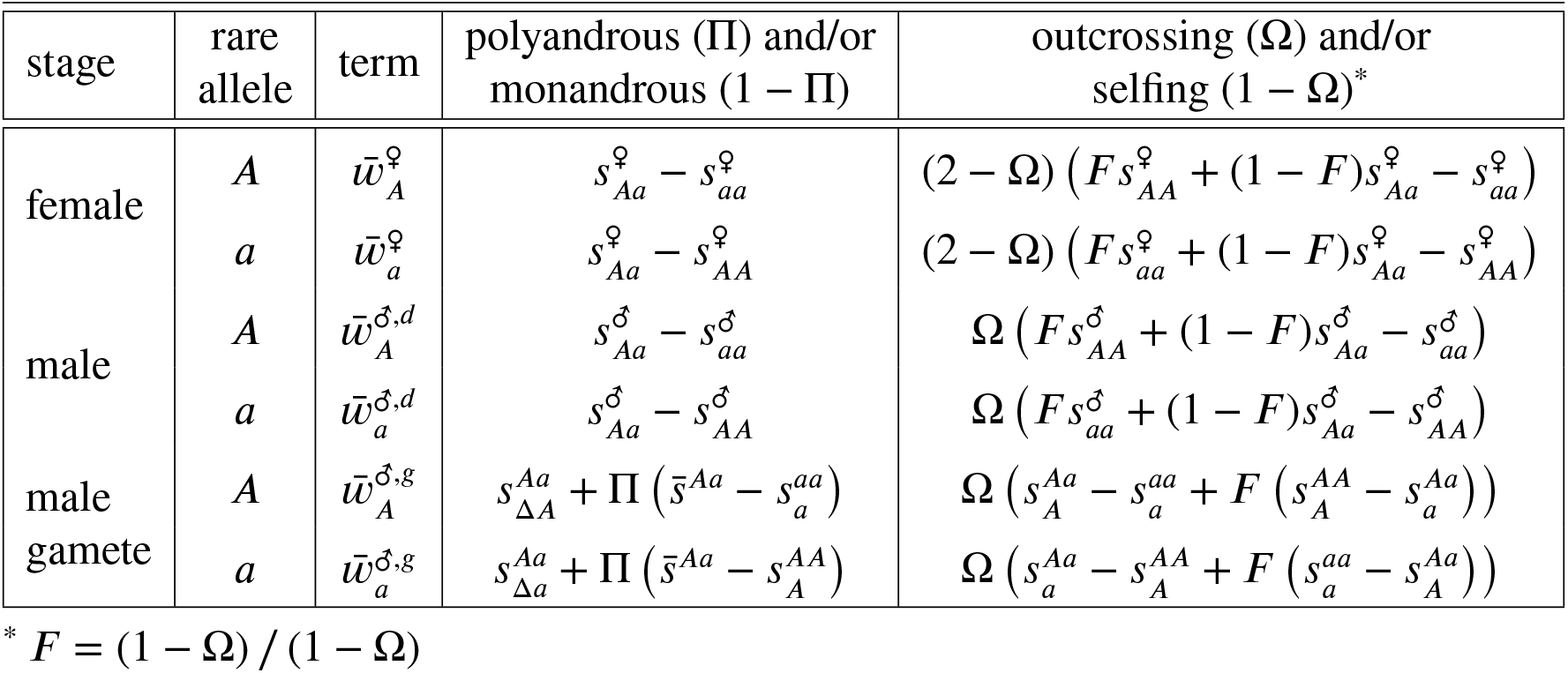
Selection terms determining invasion under different mating systems.

The selection terms in Table 2 highlight some important differences between mating systems. First, unlike monandry, selfing creates homozygotes such that homozygous fitnesses appear along with the inbreeding parameter (*F*), which indicates the excess of homozygotes relative to Hardy-Weinberg expectations. Second, male and female fitnesses are weighted equally under monandry/polyandry but unequally when there is selfing. Increased selfing (lower Ω) effectively increases the importance of female fitness and decreases the importance of male fitness (see also Jordan and Connallon 2014).

### Equilibrium Allele Frequency

Beneficial alleles will increase in frequency when rare (e.g., allele *A* when *I_A_* > 0, equation 1) and continue to be favoured when common (*I_a_* < 0), quickly spreading to fixation. However, mutational input can maintain deleterious alleles (e.g., allele *a* when *I_a_* < 0), despite selection removing them. Such a scenario of ‘mutationselection balance’ can maintain genetic variation over long time periods. The introduction of deleterious mutations decreases fitness leading to ‘mutation load’. We first calculate the expected frequency of deleterious alleles that are maintained by mutation, which can be used to calculate the per-locus mutation load.

Assuming that selection is weak and mutation rate is very small, the expected frequency of a deleterious allele at mutation selection balance is

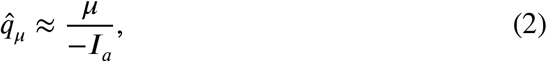

which is the ratio of the rate that new deleterious alleles arise by mutation and the rate they are removed by selection when rare.

Figure 2 shows how mating systems, gametic selection, and expression interact to affect mutation load. The degree of monandry or polyandry only affects deleterious allele frequency if there is some gametic selection (Figure 2a and Table 2). Gametic selection removes deleterious alleles and so reduces mutation load. We find that monandry increases the frequency of deleterious mutations (see Supplementary Data for proof). Monandry creates some highly competitive environments for gametes (from heterozygous males) and some highly uncompetitive environments (from homozygous males, see Figure 1b). However, the overall effect is that monandry reduces the intensity of gametic selection and increases mutation load (Figure 2a). Haploid expression further decreases mutation load by directly exposing an allele’s deleterious effects to selection. Monandry and diploid gametic expression, on the other hand, prevents gametic selection altogether because there is no fitness variation among gametes produced by a single male with diploid expression (Figure 1b).

**Figure 2:**
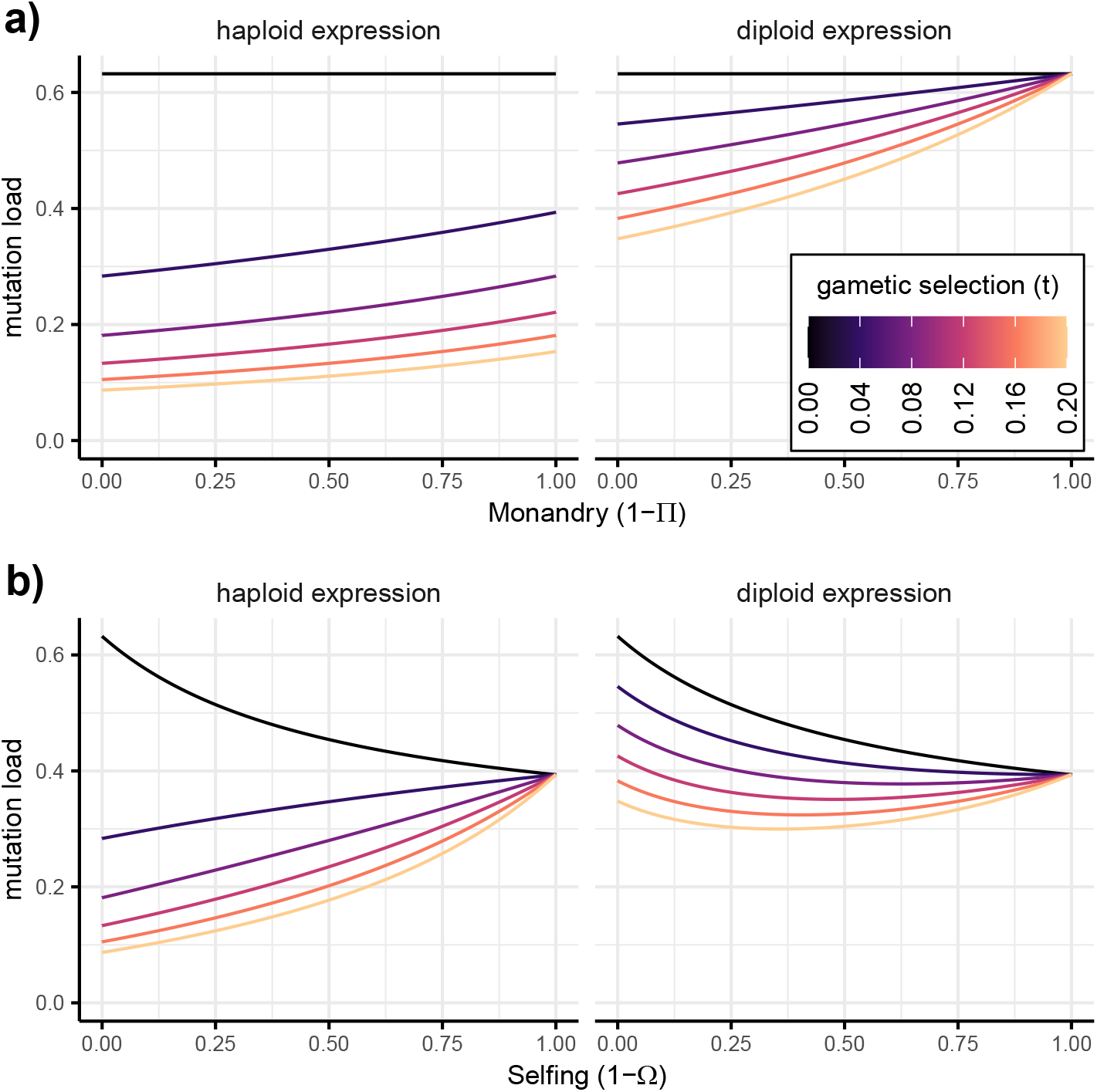
Mutation load for different mating systems and strengths of gametic selection. Mutation load is calculated for unlinked recessive deleterious alleles experiencing gametic selection (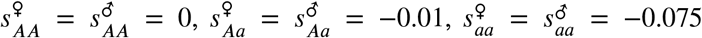, and *H* = 2), maintained by mutation at a rate of one mutation per diploid genome per generation (*U* = *2μl_μ_* = 1). The mating system varies (a) from polyandrous (Π = 1) to monandrous (Π = 0) or (b) from outcrossing (Ω = 1) to selfing (Ω = 0) and the colour shows the strength of gametic selection.

Figure 2b shows a rather different relationship between selfing and mutation load. Selfing increases homozygosity, which means that there are no heterozygotes and so no gametic selection in fully selfing populations. Selfing also exposes deleterious alleles to selection in homozygous adults, which reduces the equilibrium mutation load without gametic selection (Figure 2b). Nevertheless, when there is gametic selection, mutation load may increase with increased selfing because the intensity of gametic selection is reduced (Figure 2b).

Overall, mutation load is predicted to be lower in genes that experience gametic selection, especially if they have haploid expression. The effect of gametic selection on mutation load is eliminated under selfing or under monandry with diploid expression. Generally, we predict the difference in mutation load between genes that are involved in gametic selection and those that aren’t becomes less as monandry and selfing become common. When comparing monandrous and polyandrous populations or species, we predict increased mutation load in the monandrous populations, all else being equal. Selfing can increase or decrease mutation load relative to outcrossing. Across the genome as a whole, selfing populations are likely to have lower mutation load, but we predict this effect is less strong (or even reversed) in the subset of genes that are involved in gametic selection.

Another way that genetic variation is maintained over long time periods is via balancing selection. A classic form of balancing selection is overdominance, where heterozygotes have a higher fitness than either of the two homozygotes. Other scenarios of balancing selection include sexually antagonistic selection, where one allele increases male fitness but decreases female fitness, and ploidally antagonistic selection, where one allele increases fitness during gametic selection but decreases fitness when expressed in the diploid adults. All forms of balancing selection mean that both alleles are favoured when they are rare (*I_A_* > 0 and *I_a_* > 0). The allele frequency is then expected to reach an intermediate equilibrium and both alleles are maintained in the population. While it may be rare on a per-locus basis, balancing selection can account for an outsized fraction of genetic variation because the equilibrium allele frequencies can be high.

Under our weak selection assumptions, the equilibrium allele frequency of alleles maintained by balancing selection is

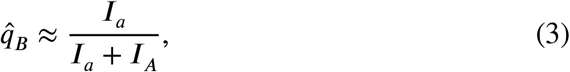

which reflects a balance between the advantage experienced by each allele when rare.

Both monandry and selfing decrease the overall intensity of gametic selection, despite creating some highly competitive environments (involving heterozygous males, Figure 1b). Thus, the allele favoured in male gametes is expected to be found at lower frequency with increasing monandry or selfing (Figure 3). Gamete-beneficial alleles are selected more strongly when they have haploid expression, which exposes alleles directly to selection, whereas diploid expression allows fitness effects to be masked. In Figure 3, we show an example of balancing selection caused by ploidally antagonistic selection, where alleles have opposite fitness effects in gametes and adults, but these conclusions also apply to other forms of balancing selection where one allele has an advantage during gametic selection.

**Figure 3:**
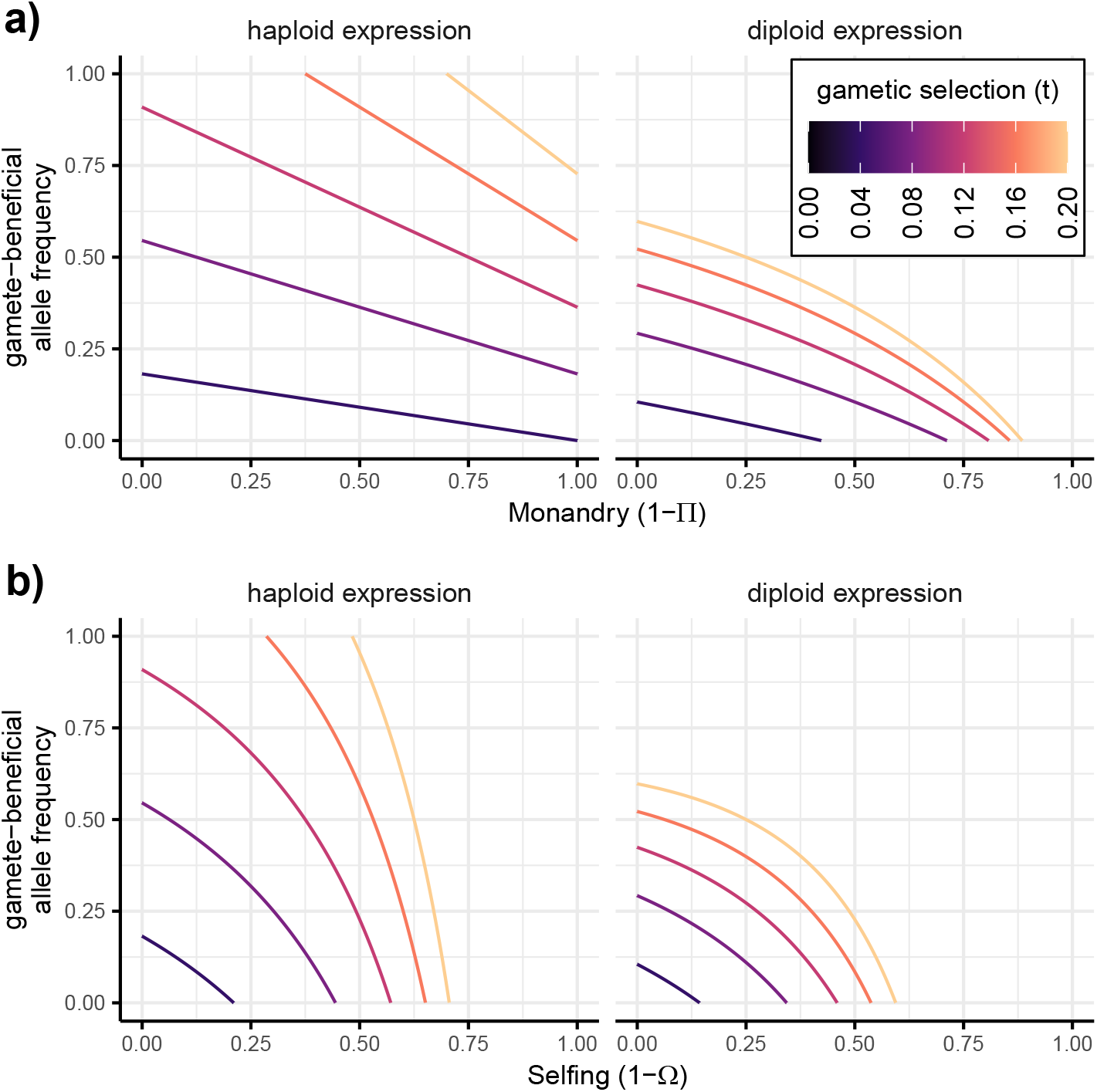
Equilibrium allele frequency under balancing selection across mating systems. We plot the equilibrium frequency of the *A* allele 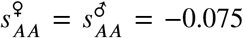, which is favoured during gametic selection. Here, selection is ploidally antagonistic because allele *a* is favoured during selection in adults 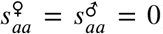. Thus, no genetic variation is maintained without gametic selection (*t* = 0). We assume beneficial effects are partially dominant (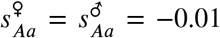 and *H* = 2). The mating system varies (a) from polyandrous (⊓ = 1) to monandrous (⊓ = 0) or (b) from outcrossing (Ω = 1) to selfing (Ω = 0) and the colour shows the strength of gametic selection (*t*).

### Mating System Evolution

We have shown that mating systems can determine what alleles are favoured, how strongly they are favoured, and shape the genetic variation that is maintained by mutations or balancing selection. Now, we explore how mating systems are expected to evolve when there is gametic selection. To examine the direction of mating system evolution, we evaluate the spread of a rare modifier allele (*m*) that changes the mating system. We assume that the modifier allele has no direct fitness effect. Therefore, there must be genetic variation at the selected **A** locus for mating system evolution to occur via its effects on gametic selection. We will assume that genetic variation is maintained by either mutation-selection balance or by balancing selection.

First, we consider modifier alleles that increase or decrease the rate of polyandry versus monandry (indicated by superscript Π). When rare (e.g., a new mutant), the *m* allele frequency changes at rate *λ*^Π^ and will increase if *λ*^Π^ > 1. Assuming that the **A** locus is at mutation-selection balance (indicated by subscript *μ*), a rare modifier allele spreads at rate

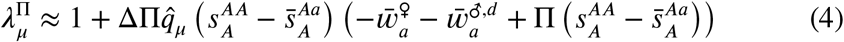

where ΔΠ = (Π*_Mm_* – Π)/2 is the increase in polyandry caused by the rare modifier allele. Because we assume that the *a* allele is deleterious at mutation-selection balance, all the other terms are positive. Thus, modifier alleles will spread 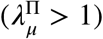 if they increase the rate of polyandry (ΔΠ > 0). That is, we expect alleles at mutation-selection balance to favour the evolution of polyandry.

We also look at the evolution of polyandry versus monandry when genetic variation is maintained by balancing selection (indicated by subscript *B*). A mutant that alters the rate of polyandry will spread if 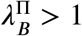 where

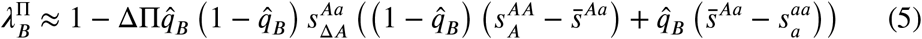

Unlike mutation-selection balance, balancing selection favours the evolution of monandry. That is, the modifier allele increases in frequency 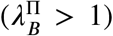 when it increases monandry (ΔΠ < 0). The other factors in equation (5), must combine to give a positive term as long as gametic fitness increases monotonically with increased expression of the higher fitness allele (e.g., when 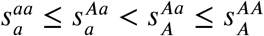, Figure 1a). The strength of selection for monandry depends on the allele frequency 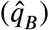 and the strength of gametic selection. However, when gametic expression is diploid, there is no fitness variation among gametes from male homozygotes 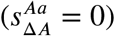 and no mating system evolution 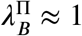 (Figure 4a).

**Figure 4:**
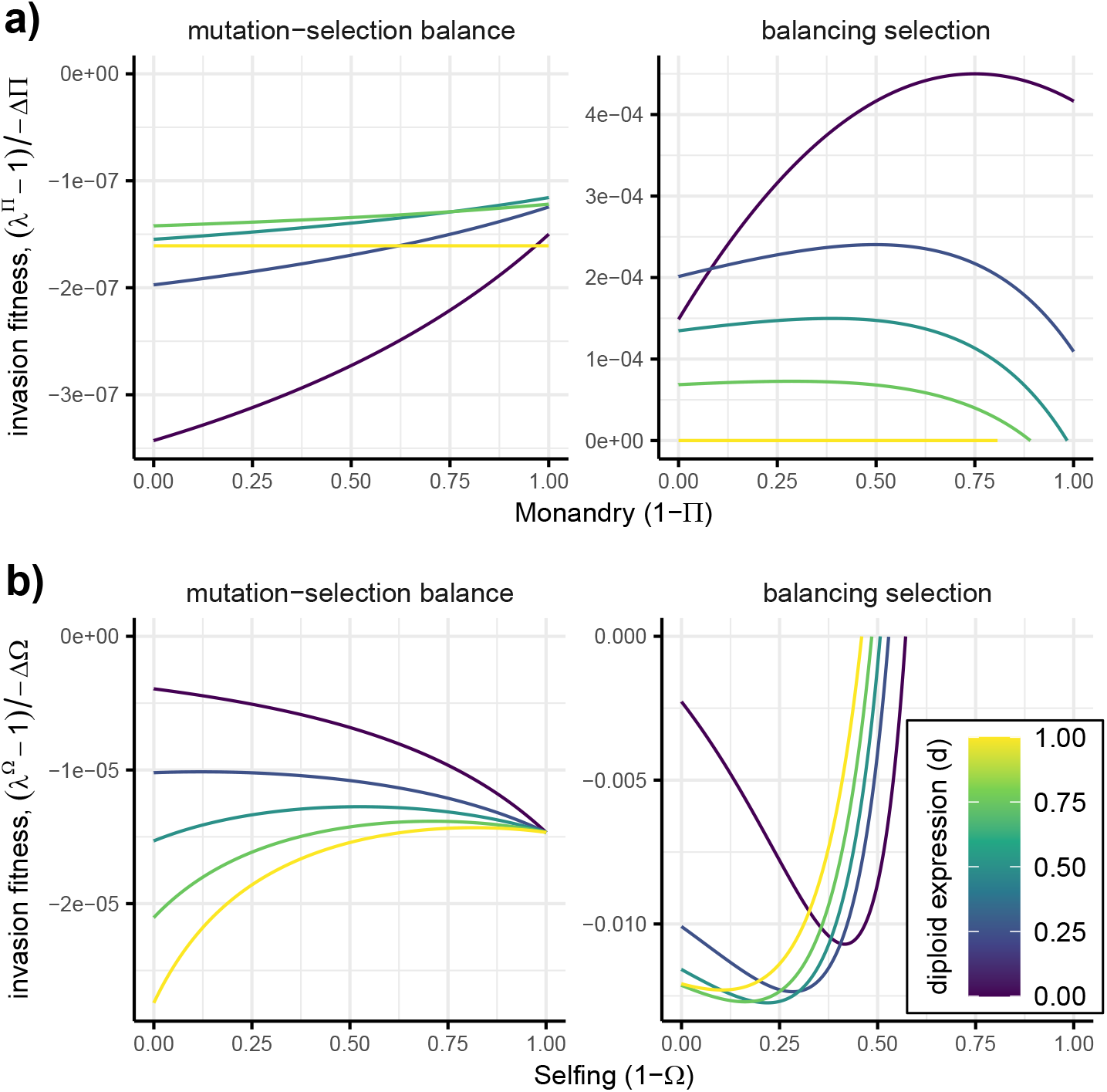
Selection for rare alleles that increase the rate of (a) monandry or (b) selfing. Negative values indicate selection for increased (a) polyandry or (b) outcrossing. Colours indicate to degree to which expression in male gametes is haploid or diploid. The *A* allele is assumed to be favoured in male gametes (*t* = 0.1). Alleles at mutation-selection balance are partially recessive (other parameters as in Figure 2) and balancing selection is ploidally antagonistic (other parameters as in Figure 3). We assume complete pollen discounting (*c* = 1) such that selfing does not have a direct transmission advantage.

The evolution of polyandry or monandry (equations 4 and 5 and Figure 4a) can be interpreted as ways to increase offspring fitness through postcopulatory sexual selection. In the case of deleterious alleles maintained at mutation-selection balance, gametic selection increases offspring fitness by removing alleles that reduce fitness in both gametes and adults. Polyandry increases the efficacy of gametic selection (e.g., Figure 2 and Figure 3). Therefore, offspring fitness is increased by evolving polyandry because it makes gametic selection more efficient.

Balancing selection on the other hand, favours the evolution of monandry. With balancing selection, gametic selection moves the equilibrium allele frequency away from optimum for adults. This is clearly true when different alleles are favoured in gametes and adults (ploidally antagonistic selection), but also true for other forms of balancing selection, such as overdominance or sexually antagonistic selection. The result is that offspring fitness is increased by reducing the strength of gametic selection, which can be achieved by evolving monandry. Notably, mating system evolution is approximately neutral with diploid-like gametic expression because the gametic fitnesses become an extension of adult male fitness so gamete-beneficial alleles effectively benefit male offspring.

We next consider modifier alleles that increase or decrease the rate of outcrossing versus selfing (indicated by superscript Ω). To leading order, the evolution of outcrossing versus selfing is dominated by the direct transmission advantage of selfing. Specifically, to leading order, a rare modifier changes frequency at rate

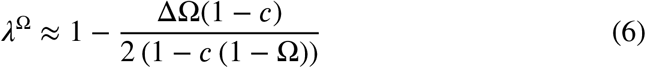

where ΔΩ = Ω*_Mm_* – Ω + *F* (Ω*_mm_* – Ω*_Mm_*) is the increase in outcrossing caused by the modifier. Without pollen discounting, selfers suffer no disadvantage in fertilising eggs from other individuals in outcrossing events. However, they monopolise the maternal and paternal contributions to zygotes formed from their own eggs, which gives selfing a strong intrinsic transmission advantage. To determine the (lower order) effect of gametic selection on the evolution of selfing, we now assume that there is complete pollen discounting (*c* = 1). That is, we assume that using male gametes for self fertilization means proportionally fewer male gametes are available to outcross and fertilize others. This eliminates the transmission advantage of selfing (equation 6) and allows us to examine the effects of selfing on offspring fitness. To simplify the results, we further assume that the modifier has a small and dominant effect on the rate of outcrossing (ΔΩ is of order *ϵ* and Ω*_mm_* = Ω*_Mm_*). When modifiers change selfing by a large amount, they can develop associations with favourable genetic backgrounds. (Campbell 1986, Charlesworth et al. 1990)

With these assumptions, a rare modifier that increases the rate of outcrossing by ΔΩ will spread at rate

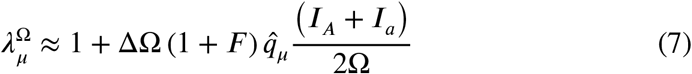

when deleterious alleles are maintained by mutation (at frequency 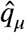, equation 2). Because the *a* allele is deleterious, *A* is favoured when rare (*I_A_* > 0) and *a* is disfavoured when rare (*I_a_* < 0), which means that selfing or outcrossing can evolve. However, most deleterious alleles are recessive, which means that the fitness difference between *AA* and *Aa* genotypes is less than the fitness difference between *Aa* and *aa* genotypes, giving *I_A_* + *I_a_* > 0. Thus, recessive deleterious alleles favour outcrossing (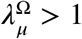 when ΔΩ > 0).

When balancing selection maintains *a* alleles (at frequency 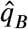, equation 3), a mating system modifier spreads at rate

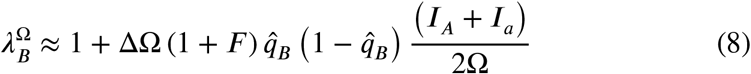

which is similar to equation (7) except 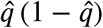 is approximately 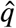 for rare alleles at mutation-selection balance. With balancing selection, both alleles have an advantage when they are rare (*I_A_* > 0 and *I_a_* > 0) and increased outcrossing is always favoured (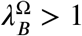 when ΔΩ *>* 0).

These results can be restated in terms of ‘inbreeding depression’ (*δ*). Inbreeding depression is calculated as 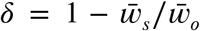 where 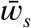 and 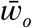 are the average fitnesses of offspring produced by selfing and outcrossing, respectively. If male gametic expression is haploid and there are no sex differences between sexes (i.e., 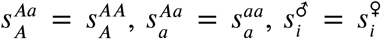), equations (7) and (8) can be rewritten as 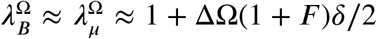. That is, inbreeding depression favours outcrossing. We have explicitly modelled the evolution of selfing in response to inbreeding depression caused by a single locus, whereas classic models (e.g., Charlesworth 1980) assume that inbreeding depression is a fixed cost.

The strongest influence on selfing evolution in our model is its intrinsic transmission advantage, which can be as strong as 50% (equation 6). Selfing also changes offspring fitness, primarily through increased homozygosity as shown by equations (7) and (8), which feature the relative fitness of heterozygotes versus homozygotes (*I_A_* and *I_a_*). Thus, despite dominating our analysis of monandry/polyandry evolution (equations 4 and 5), changes in offspring fitness via postcopulatory sexual selection have a relatively small role in selfing evolution. On a per-locus basis, the effects of selfing on offspring fitness through increased homozygosity (equations 7 and 8) are also weak but these fitness effects can combine across loci, potentially determining mating system evolution.

#### Multiple loci

A single locus has a relatively weak effect on mating system evolution. The examples in Figure 4 suggest that the selection coefficient for a modifier of monandry/polyandry is on the order of 10^-7^ when another locus is at mutation-selection balance or up to 10^-4^ when another locus experiences balancing selection. In this section, we consider the net effect of many loci on mating system evolution. First, we approximate the total strength of selection for modifiers of mating system. This selection strength can be compared against other factors in mating system evolution (e.g., cost of finding mates) to consider the relative importance of gametic selection in mating system evolution. Then, we calculate inbreeding depression across many loci, which is a crucial determinant of selfing evolution.

To consider many selected loci, we assume there is loose linkage and no epistasis so that we can ignore genetic associations between loci. This means the total indirect selection on a modifier of weak effect can be approximated by adding together the indirect selection caused by each locus, i.e., *s_net_* = *λ_net_* −1 = ∑(*λ_l_* – 1) where we sum over *l* loci to get the net selection on the mating system modifier (e.g., Otto and Bourguet 1999). We consider three types of loci: there are *l_B_* loci that experience balancing selection and *l_μ_* loci with deleterious alleles at mutationselection balance, of which a proportion *k* are expressed in gametes and (1 – *k*) only experience diploid selection. Within each of these categories, we assume that each locus is subject to the same selection coefficients for simplicity, but it is also possible to sum over a distribution of selective effects.

Figure 5 shows the net strength of selection on modifiers of monandry, which varies up to approximately 1% for these parameters. Figure 5 also shows that a relatively small number of loci under balancing selection can have a disproportionately large impact on mating system evolution. Balancing selection maintains alleles relatively high frequencies so that they can account for more genetic variation and thereby have stronger indirect effects on mating system modifiers. Because loci experiencing balancing selection favour monandry when they experience gametic selection, a small number of them could cancel out the selection for increased polyandry caused by a large number of loci at mutation-selection balance.

**Figure 5:**
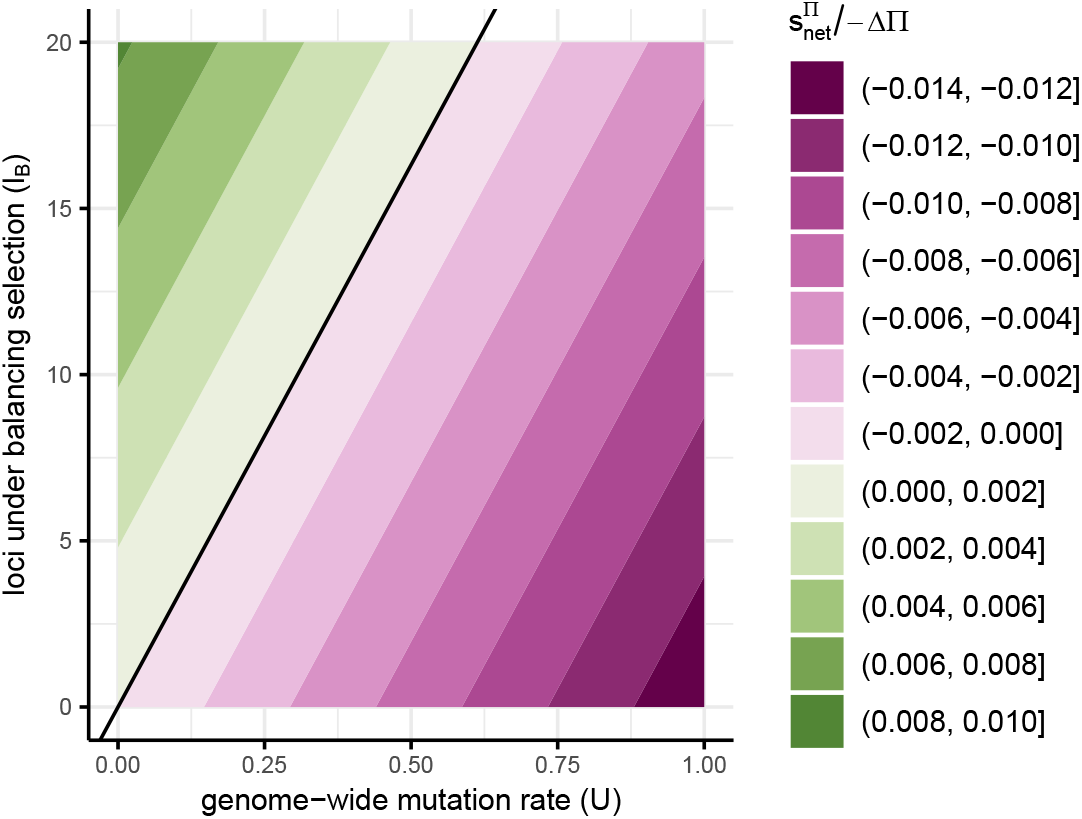
Net selection coefficient for monandry 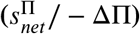 due to post-copulatory sexual selection. The diploid genome-wide deleterious mutation rate is given by twice the haploid per-locus mutation rate and the number of loci at mutation-selection balance (*U* = *2μl_μ_*). Because loci that don’t experience gametic selection have no effect on the modifier, we only include loci that experience gametic selection (*k* = 1). The strength of gametic selection is *t* = 0.12 and gametic expression is haploid-like (*d* = 0). Other parameters are the same as in Figure 2 for loci at mutation-selection balance and the same as Figure 3 for loci experiencing balancing selection.

Our modifier analysis for selfing evolution confirms that the most important factors are transmission advantage (equation 6) and inbreeding depression (equations 7 and 8), with postcopulatory sexual selection having a smaller role. The transmission advantage of selfing can be up to 50% when there is no pollen discounting. This means that outcrossing is stable when inbreeding depression diminishes the fitness of selfed offspring by more than 50% (*δ* > 0.5, e.g., equation 15 in Charlesworth 1980). The transmission advantage of selfing decreases with pollen discounting (higher *c*) such that outcrossing can be stable with less inbreeding depression. Following Charlesworth et al. (1990), we calculate the inbreeding depression across loci, which can be translated into mating system evolution for given rates of pollen discounting.

Figure 6 shows that gametic selection could have a large impact on inbreeding depression, and therefore mating system evolution. For these parameters, including gametic selection at 70% of loci decreases inbreeding depression enough to make outcrossing unstable without any pollen discounting (*δ* decreases below 0.5 between green and orange lines). This is because gametic selection can efficiently remove deleterious alleles and decrease mutation load, especially when gametic expression is haploid (Figure 2). Again, a small number of loci under balancing selection can have a disproportionately large effect on inbreeding depression because they can reach high frequencies (dashed lines in Figure 6).

**Figure 6:**
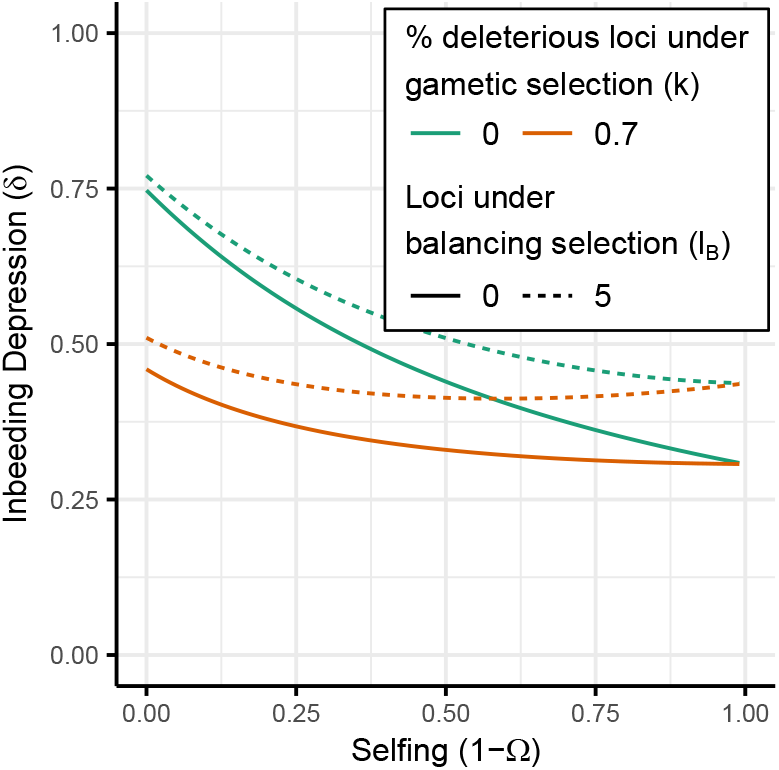
Including gametic selection at a fraction of loci (orange versus green) can significantly reduce inbreeding depression (*δ*). A small number of loci under balancing selection (dashed lines) can also have a relatively large effect on inbreeding depression. The loci under balancing selection have overdominance (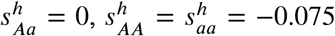, *t* = 0), which ensures the equilibrium is stable across the range of selfing rates. For loci at mutation-selection balance, the selection coefficients and genome-wide mutation rate are the same as in Figure 2 with *t* = 0.075 and haploid-like gametic expression (*d* = 0).

## Discussion

We found that monandry and selfing both decrease the efficacy of gametic selection but these mating systems follow different evolutionary trajectories. Monandry is favoured when alleles are maintained by balancing selection but not mutationselection balance (Figure 4a) whereas selfing is not favoured in either scenario unless deleterious alleles are dominant (Figure 4b). The key difference between monandry and selfing is that selfing directly increases homozygosity. The increased homozygosity caused by selfing (inbreeding depression) is more important than the effect of selfing on offspring fitness via postcopulatory sexual selection, which determines the evolution of polyandry in our model. Nevertheless, gametic selection has the potential to drastically reduce inbreeding depression by removing deleterious alleles (Figure 6), thereby causing selfing to evolve.

Despite creating some locally competitive environments (Figure 1b) selfing and monandry reduce responses to gametic selection. Responses to selection are also lessened with diploid expression due to masking effects. These patterns are in agreement with verbal arguments about the absence of pollen fitness variation under selfing (Mazer et al. 2010) and models of sperm competition that don’t include expression in other tissues (Ezawa and Innan 2013, Dapper and Wade 2016). We predict lower mutation load for genes that experience gametic selection, particularly those with haploid-expression or in polyandrous populations where gametic selection is more effective (Figure 2a). Decreasingly effective gametic selection can also mean selfing increases mutation load for genes involved in gametic selection, reversing the usual trend (Figure 2b). Overall, genes expressed in male gametes should have stronger signs of mutation load than those that aren’t but the difference between these gene sets should be less with diploid gametic expression, monandry, or selfing.

Empirical results demonstrate the expected differences in evolutionary rates for genes involved in gametic selection under outcrossing or selfing. Pollen-specific genes in outcrossing *Capsella grandiflora* had strong signatures of selection compared with seedling-specific genes (Arunkumar et al. 2013), whereas no difference was found in a predominantly-selfing species, *Arabidopsis thaliana*, after accounting for tissue specificity (Harrison et al. 2019), despite earlier reports, (Gossmann et al. 2014). Across *Arabis alpina* populations, signatures of purifying selection on pollen-expressed genes are stronger where there is more outcrossing (Gutiérrez-Valencia et al. 2022). There is also evidence that pollen phenotypes respond to mating system. Pollen tube growth rates are higher in the predominantly outcrossing *Clarkia unguiculata* than in the closely related selfing species *C. exilis* (Mazer et al. 2018) and experimental manipulation of polyandry through increasing *Mercurialis annua* plant density led to the evolution of faster pollen tube growth rates (Tonnabel et al. 2022).

Because gametic selection becomes more effective, positive and purifying selection in sperm-expressed genes should be particularly strong in genes with more haploid-like expression and in populations or species with more polyandry. These comparative predictions are valid whether genes are expressed in other tissues or not (Dapper and Wade 2016). However, we note that sperm-specific genes only experience selection in competitive sperm pools and therefore exhibit slower evolutionary responses than genes expressed in both males and females for a given selection coefficient (Dapper and Wade 2020). As expected with intense sexual selection or relaxed selection, sperm proteins show particularly rapid evolutionary rates (reviewed in Clark et al. 2006, Turner and Hoekstra 2008, Dapper and Wade 2020). Recent results from single-cell transcriptomics enable evolutionary rates to be compared between genes with different expression patterns in sperm (Bhutani et al. 2021). Indeed, due to reduced pleiotropic constraints and haploid selection, genes expressed during late stages of spermatogenesis evolve particularly rapidly (Murat et al. 2022). Extending these analyses to compare across the evolution of testes-expressed genes with haploid- or diploid-expression across different mating systems offers an opportunity to test our predictions.

As well as evolutionary responses under different mating systems, we modelled the evolution of mating systems. In general, our modifier analysis shows that mating system evolution doesn’t always maximise population-level fitness (Charlesworth et al. 1990). For example, outcrossing can increase mutation load (Figure 2b) but is still favoured when deleterious alleles are recessive. Similarly, polyandry is approximately neutral with diploid expression and balancing selection (Figure 4a) despite increasing adult survival or fecundity (Figure 3a). We assume that females control mating system evolution and that sperm/pollen is not limiting, which means - rather than optimising population-level fitness - females generally evolve mating systems that maximise the fitness of their offspring.

We examined how polyandry evolves to manipulate gametic selection. Previous work found that females should evolve traits that increase the intensity of haploid gametic selection and thereby increase offspring fitness (Otto et al. 2015). Here, we examine female mating rates as a method for manipulating gametic selection. We find that increased offspring fitness can be achieved through increased polyandry as long as the direction of selection is the same in gametes and adults. This type of genetic variation is ephemeral or maintained by mutation. When genetic variation for gametic competitiveness is maintained by balancing selection, monandry is favoured because gametic selection moves allele frequencies away from their optimum. We have not examined the maintenance of genetic variation through migration, but we hypothesise that it can allow gametic competitiveness to be positively or negatively correlated with offspring survival such that either polyandry or monandry is favoured.

We therefore show the source of correlations between sperm competitiveness and adult viability, which is crucial in previous models of polyandry evolution (Yasui 1997). At equilibrium, we find that deleterious alleles at mutation-selection balance create the positive correlation that favours polyandry whereas balancing selection creates the opposite correlation and has an outsized effect because alleles can be maintained at high frequencies (Figure 5). Furthermore, we approximate the selective advantage of polyandry in terms of standard population genetic parameters such as the genome-wide mutation rate. For the parameters using in Figure 5, polyandry can only have a weak selective advantage of, at most, around 1%. A previous analysis concluded that the ‘sexy sperm’ effect is probably too weak to favour costly polyandry (Bocedi and Reid 2015). Our analysis also shows weak indirect benefits from ‘good sperm’ such that postcopulatory sexual selection may have a minor role in the evolution of polyandry, relative to other factors (reviewed in Arnqvist and Nilsson 2000, Jennions and Petrie 2000, Arnqvist and Rowe 2005, Kokko et al. 2006, Birkhead 2010, Pitnick and Hosken 2010, Boulton 2020).

We found that the evolution of selfing was dominated by inbreeding depression and transmission advantages (Busch and Delph 2012), rather than indirect benefits from manipulating gametic selection. Nevertheless, gametic selection can be a crucial determinant of mating system evolution. Selection on gametes or gametophytes can have a large effect on mutation load (Figure 2) and inbreeding depression (Figure 6), especially with haploid expression. In turn, this can determine whether or not outcrossing is a stable strategy. For example, without pollen discounting, Figure 6 shows outcrossing can be stable without gametic selection (solid green line) but unstable when some loci experience gametic selection (solid orange line). Whether gametic selection changes the direction of selfing evolution depends on pollen discounting, mutation rates, expression and selection (Charlesworth et al. 1990), but the influence on inbreeding depression can be large. Thus, variation in the prevalence of selfing between populations or taxonomic groups could partially reflect differences in the extent of expression and selection among gametes or gametophytes, alongside other factors such as reproductive assurance (Dornier et al. 2008, Barrett 2014).

Although they are inconspicuous life cycle stages, there is considerable potential for selection when pollen or sperm compete for fertilisation. The response to selection depends on the expression of genetic material in this competitive environment and the competitors involved, which is determined by the mating system. We have explored how organisms might evolve mating systems to optimise these evolutionary responses. Predicting variation in evolutionary responses under different mating systems and expression patterns offers a way to test our understanding of evolutionary processes more generally.

## Acknowledgements

MFS is supported by a Leverhulme Trust Early Career Fellowship (ECF-2020-095). SI is supported by funding from the Natural Environment Research Council (NE/S011188/1) and the European Research Council (SELECTHAPLOID - 101001341). We thank Aneil Agrawal and Stephen Wright for helpful discussions.

## Appendix I: Two-Locus Model Details

In order to track mating events between diploids, we census the zygotic genotype frequencies (*x_ij_*) at the **A** and **M** loci (*i,j* ∈ {*MA, Ma, mA, ma*}). We assume that maternal *i* and paternal *j* haplotypes are interchangeable resulting in a total of ten diploid genotypes, *ij*. To simplify notation, we use functions *f*(*ij*) ∈ {*MM, Mm, mm*} to denote the **M** locus genotype and *g*(*ij*) ∈ {*AA, Aa, aa*} to give the **A** locus genotype for *ij* diploids. These are simply indicator functions that give the one locus genotypes from both two-locus haplotypes. In species with separate sexes, we assume that the probability of developing as a male (***♂***) or female (***♁***) is independent of the genotype at the **A** and **M** loci. After selection, genotype frequencies for adults of sex *h* ∈ {*♂, ♁*} are given by

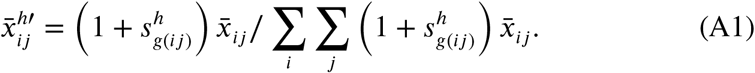

After selection among diploid adults, gametes are produced. Before mutation and selection, the frequency of gametes of genotype *k* ∈ {*MA, Ma, mA, ma*} produced by adults of sex *h* with genotype *ij* is given by 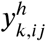. Gametes inherit parental haplotypes (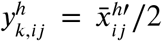 for *k* = *i* and *k* = *j*), unless the parent is a double heterozygote (e.g., *i* = *MA* and *j* = *ma* or *i* = *Ma* and *j* = *mA*). In double heterozygotes, we have to take account of the recombination rate between loci and 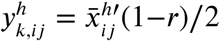 when *k* = *i* or *k* = *j* and 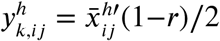 otherwise. No other gamete genotypes are possible, 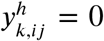 for all other combinations of *i, j, k*. We then assume that mutation from *A* to *a* occurs at rate *μ*. That is, after mutation, the gamete/gametophyte frequencies are 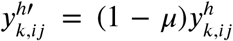 when *k* = *MA* or *k* = *mA* and 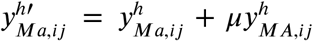. and 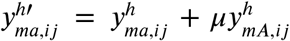. In short, we calculate the gamete/gametophyte genotype frequencies produced by adults of different genotypes after diploid selection and meiosis (with recombination and mutation).

We assume that male gametic fitness depends on the expression of *A* versus *a* alleles. That is, *A*-bearing gametes from *Aa* males can express their own genotype (100% *A* alleles), or their father’s genotype (50% *A* alleles), or some combination. The **A**-locus genotype of a gamete with haplotype *k* is indicated by *G*(*k*), where *G*(*k*) ∈ {*A, a*}, and the **A**-locus genotype of the father is given by *g*(*ij*), as above. The fitness of a male gamete with genotype *k* produced by a male with genotype *ij* is 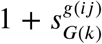. *AA* males can produce *a*-bearing gametes through mutation, but these do not feature in our results so 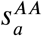 is not included in Table 1.

When we present numerical results, we must specify a continuous relationship between allelic expression levels and male gametic fitness. We assume that *AA* males produce *A*-bearing gametes with relative fitness 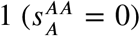 and *aa* males produce *a*-bearing gametes with fitness 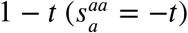. New mutatations in *AA* males can also produce *a*-bearing gametes with fitness 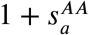, but these are rare and do not feature in our results. The fitness of gametes from *Aa* males depends on whether gametic expression is haploid (*d* = 0) or diploid (*d* = 1), where *A*-bearing gametes have fitness 1-*γ*(*d*/2)*t* and *a*-bearing gametes have fitness 1-*γ*(1-*d*/2)*t* As *a*-allele expression (*x*) increases, *γ*(*x*) increases continuously from 0 to 1. We assume *γ*(*x*) = 1 – (1 – *x^H^*)^1/*H*^ such that the *A* allele is dominant when *H* > 1 and recessive when 0 < *H* < 1 (assuming 0 < *t* < 1 such that the *a* allele is deleterious).

Male gametes compete under the mating system that is specified by the **M** locus. Under monandry or selfing, male gametes produced by a single individual compete with one another for fertilisation. In these matings that involve one male/hermaphrodite of genotype *ij*, the male gamete allele frequencies after haploid selection are

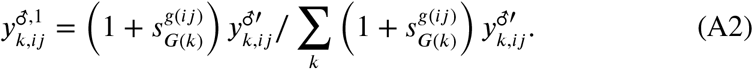

Under polyandry or outcrossing, all male gametes compete in a common pool such that, after haploid selection, the frequency of male gametes with genotype *k* is

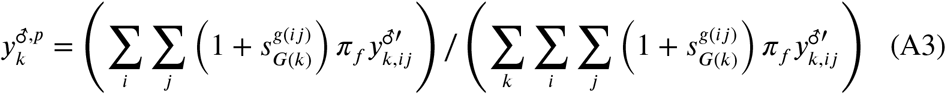

where *π_f_* = 1 – *c* (1 – Ω*_f_*) accounts for ‘pollen discounting’ via parameter *c*. When *c* = 1, selfing results in a proportional decrease in the number of male gametes that are available for outcrossing. When *c* = 0, individuals donate the same number of male gametes to a common gamete pool for outcrossing, irrespective of their selfing rate. For polyandry, we assume that *c* = 0 because there is no selfing.

First, we assume that the **M** locus controls the degree of monandry/polyandry. Specifically, a fraction Π*_f_* of the eggs/ovules produced by a mother with genotype *f*(*ij*) at the **M** locus is mated polyandrously and the remaining fraction (1 – Π_*f*(*ij*)_ is mated monandrously. Thus, we consider one hundred possible mating combinations between the ten female and male genotypes to get the zygotic genotype frequencies in the next generation, given by

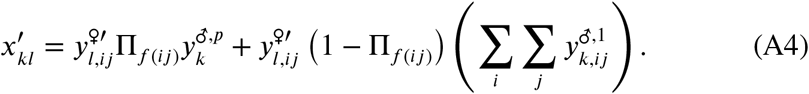

We assume that the choice of mate is random with respect to the **A** and **M** loci. Thus, we only model sexual selection that is post-copulatory and caused by gametic selection.

Second, we assume that the **M** locus controls the degree of selfing versus outcrossing. Specifically, a fraction of eggs/ovules are specified to mate via outcrossing, Ω_*f*(*ij*)_, or selfing, (1 – Ω_*f*(*ij*)_). For example, a fraction (1 – Ω_*f*(*ij*)_) of flowers may remain closed (cleistogamous) and self fertilise while the other flowers open (chastogamous) and outcross. The zygotic genotype frequencies in the next generation are given by

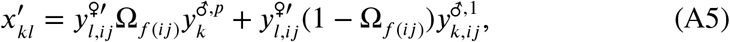

such that offspring produced by selfing are derived from the same individual with the same genotype. In equations (A4) and (A5), the paternally-inherited haplotype has index *k* and the maternally-inherited haplotype has index *l*. These are equivalent and are combined to get the ten zygotic allele frequencies for the next generation.

Note that gametic selection under selfing or monandry is determined by the same equation (A2), which sums over *k* gamete genotypes produced by a single male. Polyandrous matings and outcrossing allow competition between gametes from multiple males so equation (A3) also includes summations over all *ij* male genotypes. Under monandry, a female can mate with any male so equation (A4) sums over all *ij* male genotypes. On the other hand, the mother and father is the same when selfing, equation (A5).

## Appendix II: Mutation Load and Inbreeding Depression

We calculate mutation load and inbreeding depression based on the equilibrium allele frequency for single loci presented in the main text. Mutation load is the reduction in mean fitness caused by deleterious mutations. Inbreeding depression, *δ*, is the relative reduction in fitness caused by selfing versus outcrossing, 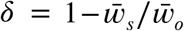 where 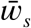 and 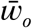 are the mean fitnesses of offspring produced by selfing and outcrossing, respectively.

We consider two sets of loci: *l_B_* loci experience balancing selection and *l_μ_* loci experience purifying selection (i.e., expected to be at mutation-selection balance). Following model II of Charlesworth and Charlesworth (1992), a fraction *k* of the *l_μ_* loci experience purifying selection in both the diploid and male gametic phases while the remaining fraction are only expressed in diploid adults. The mean fitness across all loci is given by 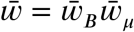 where 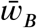 and 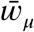 are the mean fitness effects of loci experiencing balancing selection and purifying selection, respectively.

We begin by assuming that there are no sex differences in selection and that loci of the same type have uniform effects. That is, 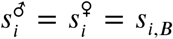 for loci under balancing selection and 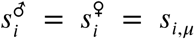 for loci at mutation-selection balance. Given that loci are unlinked and have multiplicative fitness, the mean fitness effect of loci that experience balancing selection is given by

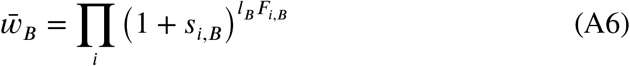

where *F_i,B_* is the equilibrium frequency of genotype *i*. The mean fitness effect of loci at mutation-selection balance is given by

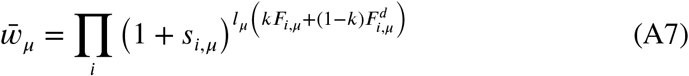

where the frequency of genotype *i* at loci with and without gametic selection is given by *F_i, μ_* and 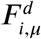, respectively. This calculation assumes that the loci that experience gametic selection have the same fitness effect on diploid adults as those that don’t experience gametic selection but this assumption can be relaxed.

Following the method of Charlesworth and Charlesworth (1992), we can approximate mutation load 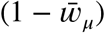 by taking sequential Taylor series for weak selection and then small mutation rates. This yields

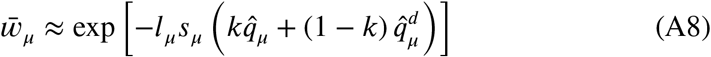

where

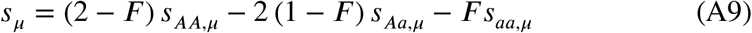

gives the diploid fitness effect of deleterious alleles, 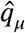 and 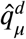 are equilibrium allele frequencies for loci that experience gametic selection (equation 2) and those that don’t (equation 2 with 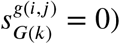).

Inbreeding depression is given by

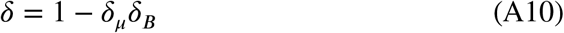

where the contributions of loci at mutation-selection balance and under balancing selection are

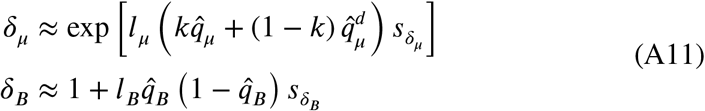

and the average diploid fitness effect of increased homozygosity is given by

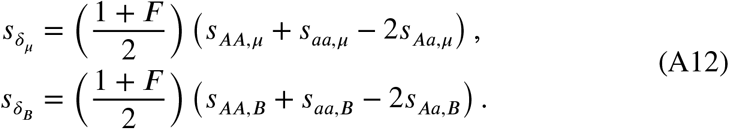

With haploid gametic expression 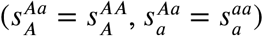 and no sex differences in selection, the growth rate of a selfing rate modifier given by equations (7) and (8) can be rewritten as 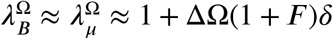, where inbreeding depression, *δ*, is calculated from a single locus. That is, with *k* =1, *l_μ_* = 1, and *l_B_* = 0

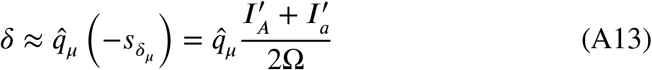

and with *l_μ_* = 0, and *l_B_* = 1

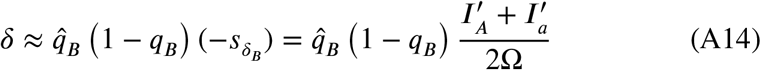

where 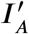 and 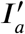 are the invasion conditions from equation (1) with haploid gametic expression and no sex differences in selection (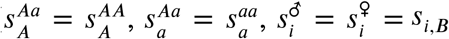, and 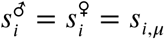).

If there are sex differences in selection and/or non-haploid gametic expression, we can interpret the modifier growth rates, 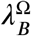 and 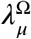, in terms of a modified version of inbreeding depression, *δ\**. To do so, we introduce a new mean fitness 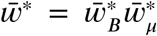 that includes gametic fitness and weights male and female fitness terms according to the outcrossing rate, as in Table 2, such that

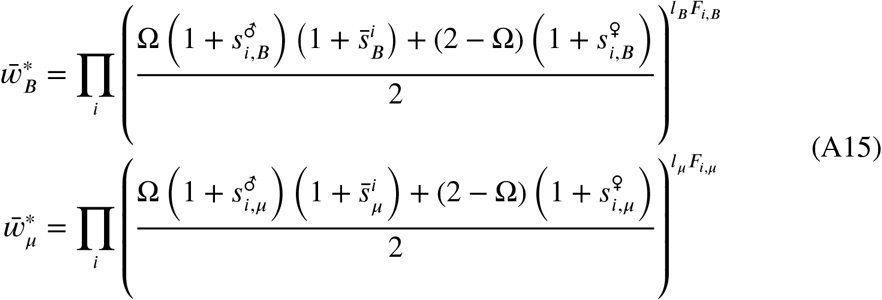

where the average fitness of gametes produced by males with genotype *i* is given by 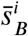 and 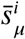 where 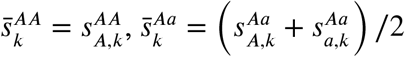, and 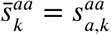.

Calculating a modified form of inbreeding depression, *δ*\*, using the weighted mean fitnesses given by equation (A15) gives

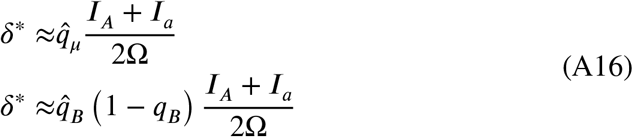

for a single locus at mutation-selection balance (*l_μ_* = 1, *k* = 1, *l_B_* = 0) or balancing selection (*ļ_μ_* = 0, *l_B_* = 1), respectively. Thus, 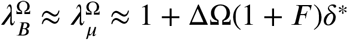 and a modified inbreeding depression that includes male gametic fitness and is weighted by the outcrossing rate gives a possible way of interpreting the growth rate of rare modifiers.

